# Identification of two principal amyloid-driving segments in variable domains of Ig light chains in AL amyloidosis

**DOI:** 10.1101/354571

**Authors:** Boris Brumshtein, Shannon R. Esswein, Michael R. Sawaya, Alan T. Ly, Meytal Landau, David S. Eisenberg

## Abstract

Systemic light chain amyloidosis (AL) is a disease caused by overexpression of monoclonal immunoglobulin light chains that form pathogenic amyloid fibrils. These amyloid fibrils deposit in tissues and cause organ failure. Proteins form amyloid fibrils when they partly or fully unfold and expose segments capable of stacking into β-sheets that pair forming a tight, dehydrated interface. These structures, termed steric zippers, constitute the spines of amyloid fibrils. Here, we identify segments within the variable domains of Ig light chains that drive the assembly of amyloid fibrils in AL. We demonstrate there are at least two such segments. Each one can drive amyloid fibril assembly independently of the other. Thus these two segments are therapeutic targets. In addition to elucidating the molecular pathogenesis of AL, these findings also provide an experimental approach to identify segments that drive fibril formation in other amyloid diseases.

---

Systemic light chain amyloidosis (AL) is a lethal disease caused by excess immunoglobulin light chains(1). Each year, approximately 4,000 patients in the U.S. and 1,500 in the U.K. are diagnosed with AL. However, recent reports indicate systemic amyloidosis is underdiagnosed(2–4). AL is frequently but not always associated with multiple myeloma(5–8). In AL, plasma cells produce excess monoclonal full-length light chains, or their variable domain segments, which circulate in the blood and assemble into pathologic amyloid fibrils. Although the sequences of light chains vary among patients, the molecular progression of the disease is similar: insoluble and degradation-resistant amyloid fibrils deposit in essential tissues and cause organ failure(9–11).

AL amyloid fibrils contain full-length light chains or just their variable domains. Overexpressed light chains (LCs) form dimers of identical LCs rather than the normal pairing of LCs with Ig heavy chains. LCs consist of two domains, variable (VL) and constant (CL), which are covalently connected by a joining segment (J). The amino acid sequences of LCs are determined by somatic gene recombination and are of two classes, lambda or kappa(12–14). In AL, the amyloid fibrils include either full-length LCs (VL-J-CL) or VLs, yet the ubiquitous presence of VLs indicates that this domain may be the minimal and essential unit for fibril assembly(15–17).

Amyloid fibrils exhibit common biochemical and structural properties. They are self-propagating, insoluble in aqueous solutions, unusually resistant to degradation, and bind the fluorescent dye Thioflavin T (ThT). Molecular structures of amyloid fibrils reveal pairs of tightly mated beta sheets, termed “steric zippers Upon exposure to X-rays, the spine gives rise to a cross-beta diffraction pattern(8, 9, 18, 19). The interdigitating side chains of segments form mating sheets with a dry, highly complementary interface, which serves as a scaffold for fibril extension(20). Once a steric zipper has formed, hydrogen bonds between the main chains propagate the spine of amyloid fibrils, while the dry interface contributes to adhesion of the mating sheets.

Proteins contain putative steric-zipper forming segments. The propensity of a segment to form a steric zipper can be estimated from its sequence(21). In globular proteins, segments predicted to form steric zippers are often buried in the interior and protected from forming fibrils. However, upon full or partial unfolding of the protein, these segments become exposed to solvent and can stack to form the amyloid fibril spine. Proteins known to assemble into amyloid fibrils may contain multiple segments capable of forming steric zippers. In the absence of experiments, it is unclear whether every, some, or only one of these segments is responsible for driving fibril formation. A protein containing more than one segment, which independently drive fibril formation, can form amyloid fibrils having different spines and thus different structures (Figure 1). The different fibril structures formed by the same protein are known as amyloid polymorphs(22–25). .

**Figure 1.**
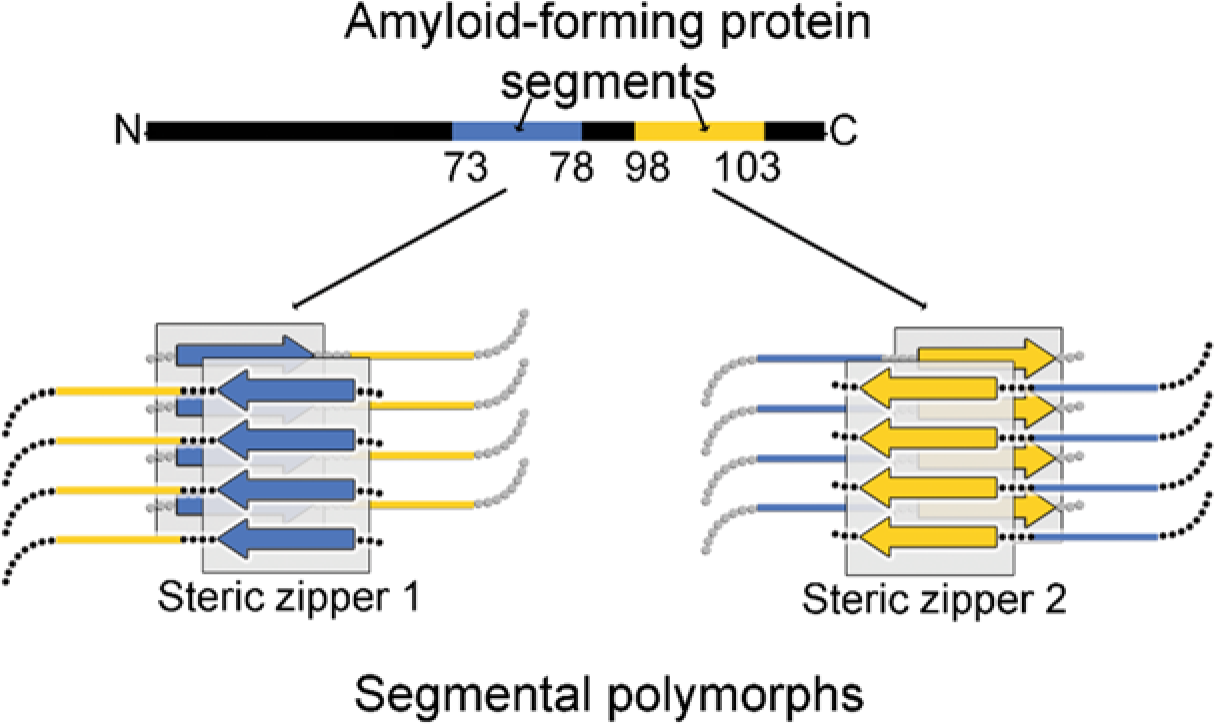
Segmental polymorphism. Amyloid fibrils form through adhesive segments that stack to form steric zippers. Ig VLs contain two segments capable of forming steric zipper fibril spines, shown here schematically in blue and yellow. Each one of the steric zipper-forming segments can induce assembly of a particular amyloid polymorph with its distinctive spine.

Here we seek the molecular basis of the formation of amyloid fibrils from Ig light chains of both lambda and kappa families. The Human Gene Nomenclature Committee lists 33 functional genes for lambda and 38 for kappa variable domains(26–28). The repertoire of VLs is greatly expanded by the linkage of lambda and kappa domains to a variety of J-segments and constant domains. Consequently the Amyloid Light Chain Database lists 616 lambda and 192 kappa VL s(17). Sequence alignments of these VLs involved in AL do not reveal a single residue or segment that exclusively accounts for their amyloidogenic property, and differences in amino acids between VLs within the same family, lambda or kappa, affect the propensity to form amyloid fibrils(29–31). Hence the identification of the amyloidogenic segments of VLs is challenging.

We define an amyloidogenic segment by two criteria: (1) introduction of amyloid-inhibiting residues, such as proline, into the segment by site-directed mutagenesis halts fibril formation of the parent VL, and (2) the segment in isolation from the remainder of the VL forms a steric zipper.

Our search for amyloidogenic segments focuses on a human genomic reference VL sequence and two patient-derived VL sequences, one kappa and one lambda. We found two amyloidogenic segments in VLs and verified that both fulfill criterion 1 by site-directed mutagenesis. Each of the two amyloidogenic segments independently causes amyloid fibril formation. Study of the isolated peptides shows that each segment also fulfills criterion 2 because each can form a steric zipper. By sequence comparison we find that homologs of these segments are amyloidogenic in both lambda and kappa families of LCs and are the key propagators of fibril formation, thereby generalizing our model of amyloid formation to other LCs and suggesting these two LC segments as therapeutic targets.

## Results

### Finding amyloid-driving segments

To identify the amyloidogenic steric zippers within VLs, we employed two complementary approaches: a computational assessment using ZipperDB to identify candidate segments with a high propensity to form steric zippers, and experimental mutagenesis to inhibit the ability of candidates to form amyloid fibrils, as detected by ThT assays(32). We identified a genomic variant of a lambda VL, 2-8 with J1 connecting segment (VL2-8-J1) as a useful reference model for our experiments; upon exposure to destabilizing conditions, VL2-8-J1 forms amyloid fibrils (Figure 2).

**Figure 2.**
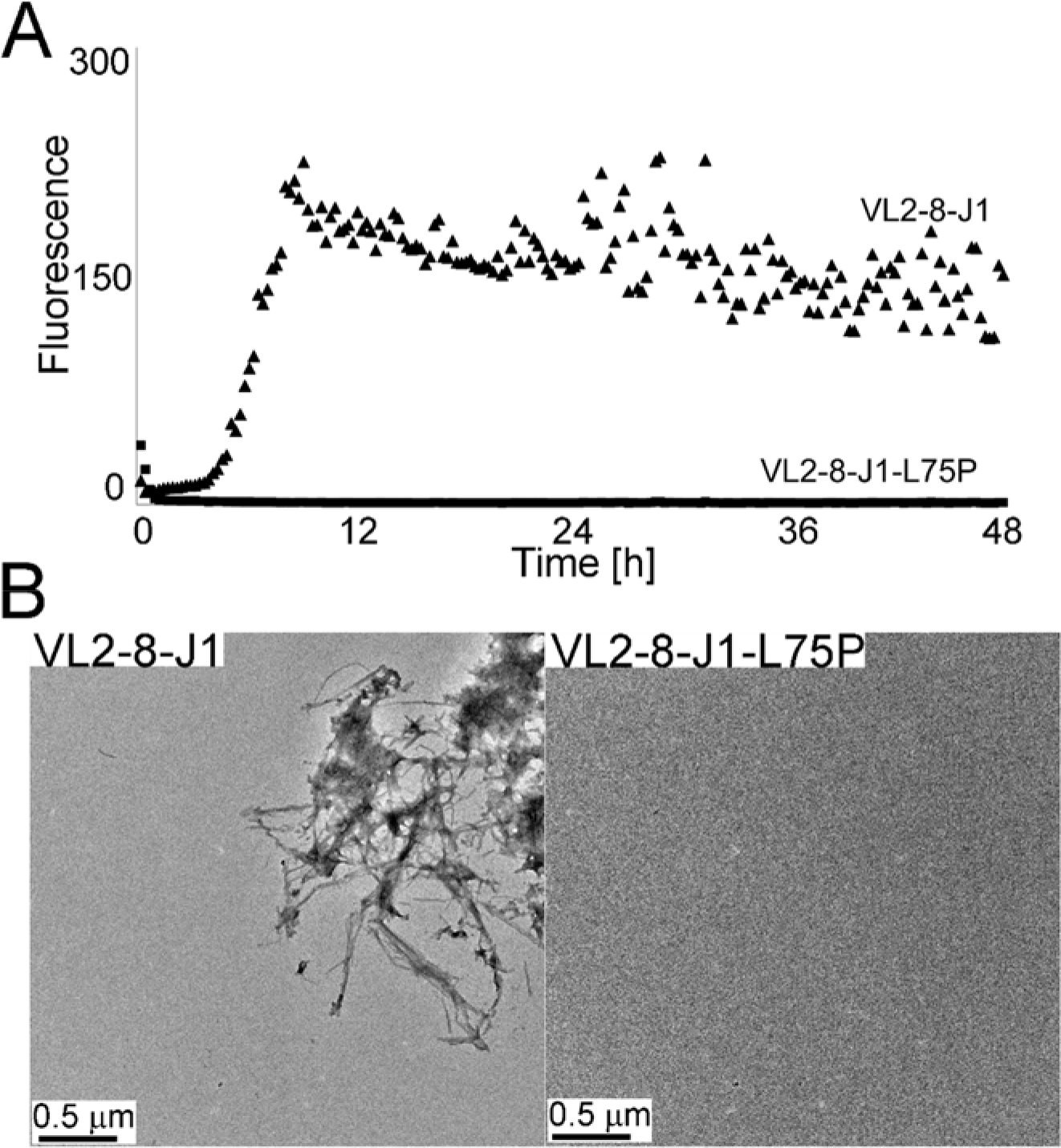
Two assays of amyloid fibril formation of the reference sequence VL2-8-J1 and its amyloid fibril-inhibiting mutation L75P: ThT fluorescence assay and electron micrographs. The ThT assay shows fluorescence, indicative of formation of amyloid fibrils (a) and electron micrographs visualize them (b). Upon exposure of VL2-8-J1 to destabilizing conditions, the protein forms amyloid fibrils. In contrast, the mutated variant, VLJ2-8-J1-L75P, under the same conditions does not show an increase in fluorescence in the presence of ThT or display amyloid fibrils in electron micrographs.

Within VL2-8-J1, ZipperDB [URL: services.mbi.ucla.edu/zipperdb/] identified five high-propensity steric zipper-forming regions, each region containing conserved residues (Figure 3, regions A-E). By using site-directed mutagenesis, we tested our predictions of involvement of these regions for formation of amyloid fibrils. Our site-directed mutagenesis replaced conserved amino acids with proline residues. Prolines introduce geometric strain into beta-sheets and eliminate a main-chain hydrogen bond, thus destabilizing the amyloid spine and impeding formation of amyloid fibrils(33). We sequentially replaced every amino acid with proline within each of the five predicted steric zipper-rich regions, A-E. A single mutation, L75P in segment C, stopped formation of amyloid fibrils by the reference sequence, VL2-8-J1, whereas prolines in other positions did not.

**Figure 3.**
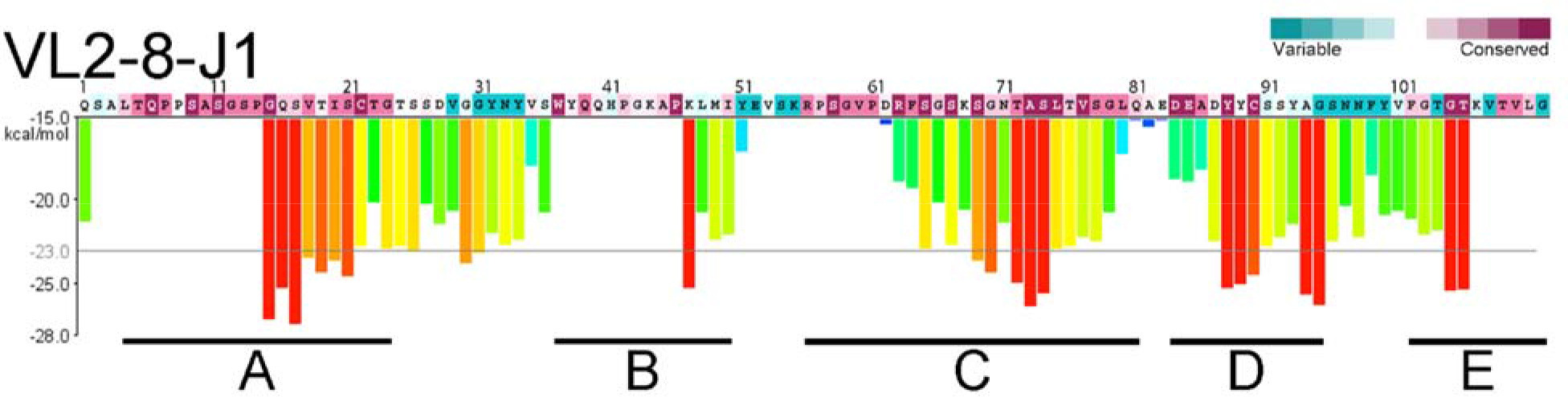
Regions predicted to contain segments capable of forming steric zippers in a genomic reference sequence VL2-8-J1. Purple and blue colors of the amino acid sequence show conserved and variable amino acids (ConSurf, http://consurf.tau.ac.il). ZipperDB (https://services.mbi.ucla.edu/zipperdb/) identified five major, high-propensity steric-zipper regions that coincide with conserved amino acids. The predicted steric zippers were the initial focus of our site-directed mutagenesis experiments. The vertical scale to the left of the histogram shows the calculated energy gain upon formation of a steric zipper. The horizontal grey line shows the energetic gain of −23 kcal/mol. Each histogram bar refers to a 6-residue segment, and orange and red bars represent segments with a high-propensity to form steric zippers. Gaps in the calculated propensities arise due to the presence of prolines, which impede ZipperDB calculations. The prediction shows 5 main regions: A includes residues 4-24, B—37-50, C—56-81, D—84-95, and E—102-111.

Upon identification of the amyloid-inhibiting proline mutation VL2-8-J1-L75P, we examined if residue L75 is required for formation of amyloid fibrils by other pathologic VLs: a lambda-type VL (Mcg) and a kappa-type VL (AL09)(34, 35). Both Mcg and AL09 are specific VL variants isolated from patients afflicted with AL. The same site-directed mutation, L75P, did not stop amyloid fibril formation by Mcg (Figure 2). The opposite effects of the L75P mutation in two different VL types provide an important insight: a single steric zipper is not responsible for formation of amyloid fibrils by all VL types. This insight led us to hypothesize that VLs may form different polymorphs, and there may be more than a single segment that can independently induce formation of amyloid fibrils.

### Finding a second amyloid-driving segment

Inability of the L75P to stop amyloid fibril formation by Mcg led us to perform a “proline-scan” experiment. With the aim of identifying segments required for amyloid fibril formation, we introduced proline mutations in 4 consecutive residues in each of 28 separate constructs that span the entire polypeptide sequence of Mcg, except that we did not replace either of the two structural cysteines with proline (Figure 4). Table 1 summarizes the results of the proline-scan experiment: all 28 mutated constructs consistently formed amyloid fibrils, except for Construct-25, which contains mutations in region E (residues F99P-V100P-F101P-G102P), in which several segments with a ZipperDB-predicted high propensity for amyloid formation are located. For this construct, several independent batches formed amyloid fibrils while others did not. These data suggest that at least for Mcg, no single steric zipper individually accounts for the amyloid-forming property of VLs, and several different polymorphs may exist. When combined, L75P and F99P-V100P-F101P-G102P mutations abolished the ability of Mcg to form amyloid fibrils (Figures 5 and 6). To summarize, every Mcg construct with tetra-proline mutations covering the entire VL sequence forms amyloid fibrils. However, the Mcg with L75P and F99P-V100P-F101P-G102P together does not form fibrils. These observations support our conjecture that no single steric zipper fully accounts for formation of amyloid fibrils by VLs; rather at least two segments drive formation of steric zippers.

**Figure 4.**
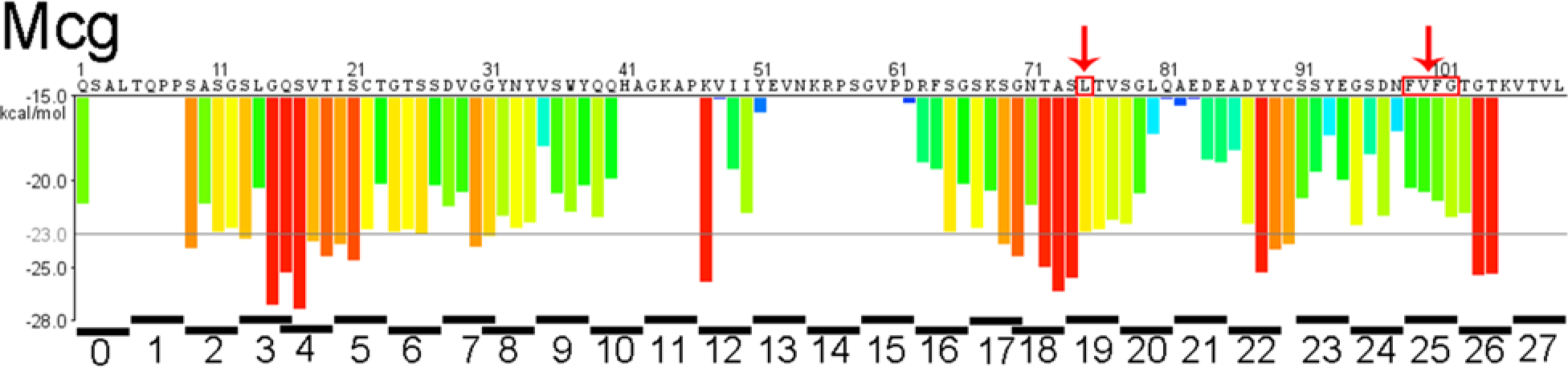
Predictions based on proline scanning of steric zipper propensity for segments of the Mcg sequence from an AL patient. The Mcg sequence is shown on top of the histogram. Four-residue segments of the proline-scanning, site-directed mutagenesis experiment are shown in black bars below the energy histogram. Peptide-scanning was performed by mutating four consecutive residues in each segment to proline. The mutated residues are identified at the bottom of the histogram and represent individual constructs each containing a tetra-proline replacement. No single tetra-proline mutation inhibited amyloid fibril formation of Mcg, but constructs L75P and 25 together did. Red arrows identify amino acids that were identified as involved in formation of amyloid fibrils.

**Table 1.** The effects of site-directed mutations on amyloid fibril formation of three different VLs. The tri-partite experiment comprises (1) an amyloid-inhibiting mutation in a reference variant of a lambda VL, (2) a lambda variant from a patient, and (3) a kappa variant from another patient. The main conclusion is that there are two amyloidogenic segments in VLs capable of driving formation of amyloid fibrils.

| Variant or mutant | Increase in ThT fluorescence | Detection of fibrils in electron micrographs | Ability of the site-directed mutation to stop amyloid fibrils |
| --- | --- | --- | --- |
| (1) A genomic lambda variable domain |  |  |  |
| VL2-8-J1 | + | + |  |
| VL2-8-J1-L75P | - | - | Positive |
| (2) A lambda variable domain from an AL patient |  |  |  |
| Mcg | + | + |  |
| Mcg-L75P | + | + | Negative (1 segment) |
| Mcg-F99-V100-F101-G102 | +/-* | +/-* | Inconclusive (2 segments) |
| Mcg-L75P-F99-V100-F101-G102 | - | - | Positive (2 segments) |
| All other constructs | + | + | Negative (2 segments) |
| (3) A kappa variable domain from an AL patient |  |  |  |
| AL09 | + | + |  |
| AL09-F73P | + | + | Negative |
| AL09-Y96P-T97P-F98P-G99P | + | + | Negative |
| AL09-F73P-Y96P-T97P-F98P-G99P | - | - | Positive |

\* "+/-" indicates that some batches formed amyloid fibrils while others did not. Three batches formed fibrils, two did not. All other mutants were tested in at least three batch-independent repetitions.

To verify whether these two segments are essential for formation of amyloid fibrils by another VL type, we introduced the corresponding mutations into a well-studied, patient-derived pathologic kappa VL, AL09(36). Sequence alignment of AL09 and Mcg identified the corresponding mutation sites: Mcg-L75P corresponds to AL09-F73P, and Mcg-F99P-V100P-F101P-G102P corresponds to AL09-Y96P-T97P-F98P-G99P. As with Mcg, only the combined mutant, AL09-F73P-Y96P-T97P-F98P-G99P, stopped amyloid fibril formation; the individual mutations did not affect the amyloidogenic potential of AL09 (Figure 5). The reproducibility of results in both types of VLs, lambda and kappa, corroborates the involvement of two segments in formation of VL amyloid fibrils.

**Figure 5.**
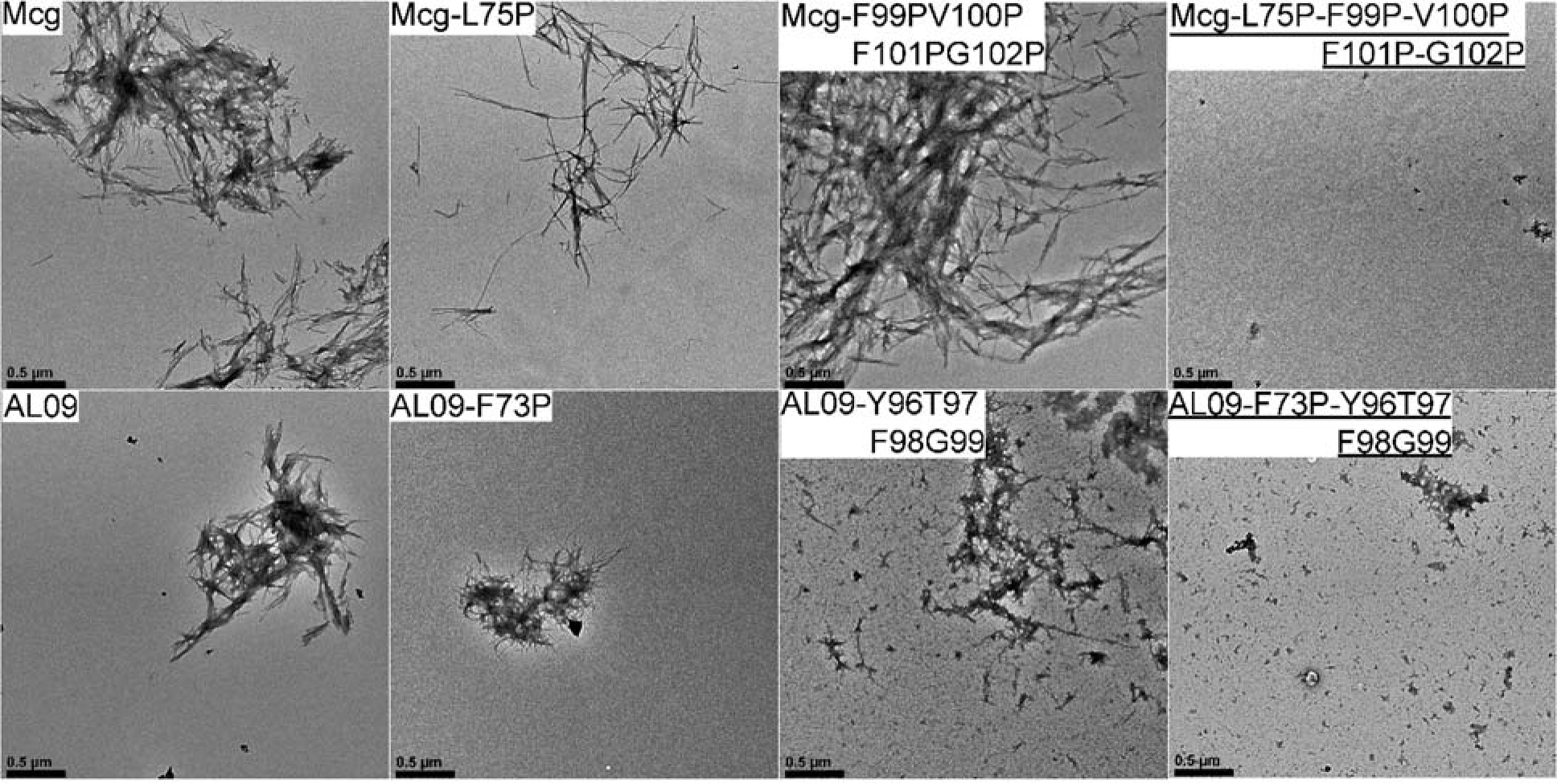
Negative-staining electron micrographs of amyloid fibrils formed by Mcg and AL09 sequences. The top row shows the effect of mutations in Mcg on the ability to form amyloid fibrils: Simultaneous mutations in two different segments abolish the ability to form amyloid fibrils. The bottom row shows the same effect of the corresponding mutations in AL09. The amyloid fibril inhibiting mutations are underlined. Black scale bars represent distances of 0.5 microns.

### Atomic structures of amyloid-driving segments

Crystal structures of individual peptides derived from amyloidogenic segments, two from the lambda VL Mcg (ASLTVS and NFVFGT) and one from the kappa AL09 (YTFGQ), reveal that these peptides form steric zippers (Figure 6). Each of these steric zippers features the typical dry interface with high-surface complementarity. The crystals of the fourth segment, EFTFTIS from kappa AL09, were of insufficient resolution to resolve its atomic structure. However, cylindrical averaging of the single-crystal diffraction data shows the characteristic reflections of steric zippers (Figure 10), as do the fibril diffraction patterns calculated from the atomic coordinates of the other three.

**Figure 6.**
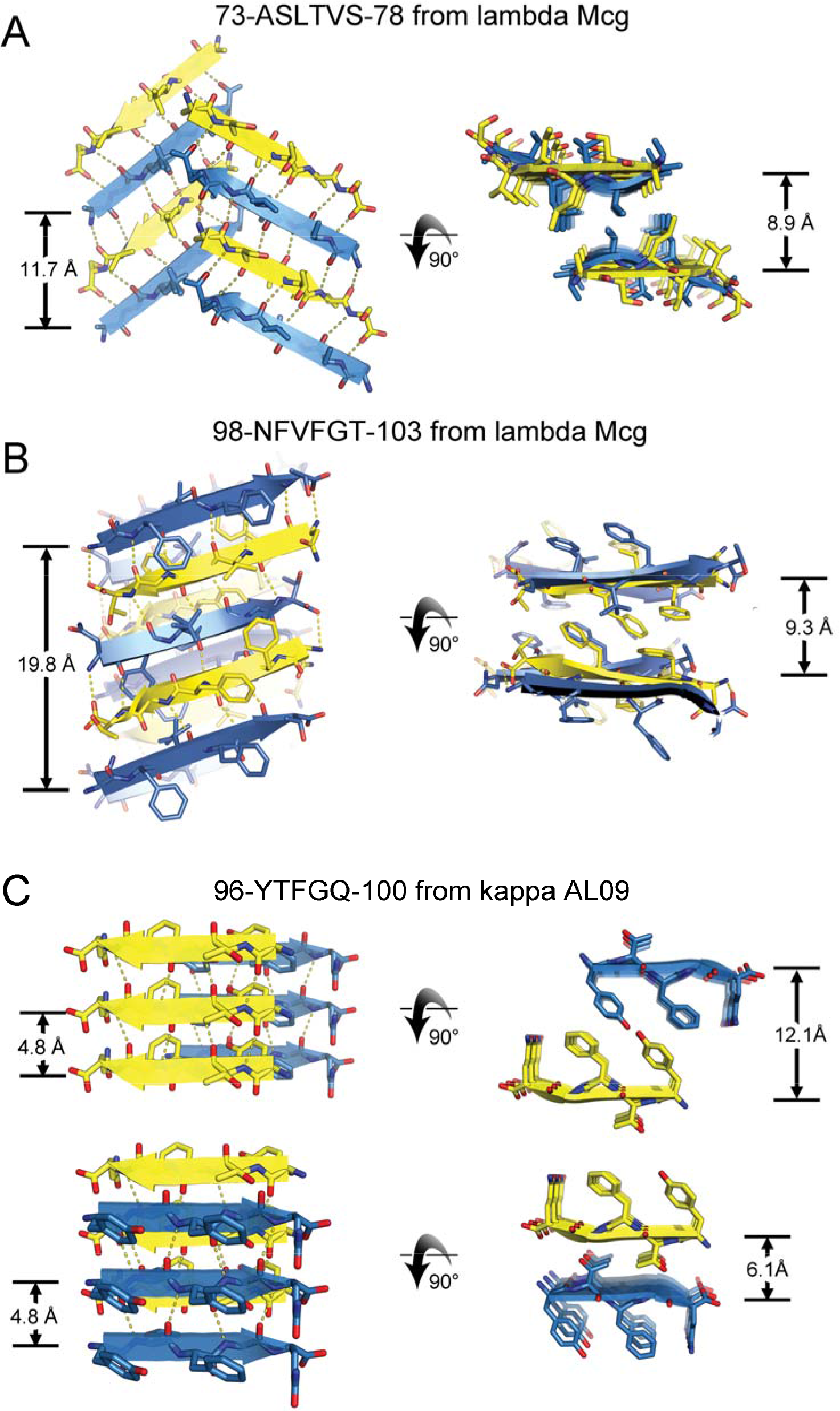
Crystal structures of VL segments that drive the assembly of amyloid fibrils. Each of the three segments adopted a steric zipper motif, characterized by pairs of sheets, tightly mated by the interdigitation of side chains across the interface and exclusion of water molecules. For each segment, two views are given: fibril axis vertical (left column) and fibril axis normal to the page (right column). Sheet architecture and symmetry differ among the three segments as follows. **A.** Segment 1 of Mcg, ASLTVS. Sheet architecture is anti-parallel, antifacial, and out-of-register (Class 5). The translational repeat distance along the fibril axis is 11.7 Å. **B.** Segment 2 of Mcg, NFVFGT. Each sheet contains four orientations of strands in the asymmetric unit, creating a level of complexity beyond any of the ten previously described steric zipper symmetry classes. The sheet is out-of-register, with an unusually long translation repeat distance of 19.8 Å. **C.** Segment 2 of AL09, YTFGQ. Sheet architecture is parallel, in-register, with a conventional translational repeat distance of 4.8 Å. Two types of dry interfaces are evident in the crystal packing; face-to-face, and back-to-back (both Class 1). Distances between sheets are given in the right column.

Our three crystal structures of VL amyloidogenic segments reveal steric zippers belonging to different symmetry classes(20). Symmetry classes define the spatial relationship between identical strands in the zipper. The steric zipper of segment YTFGQ from AL09 is composed of pairs of parallel sheets mated face-to-face, identifying it as Class 1 (Figure 6C). The same crystal structure also reveals a different pair of sheets mated back-to-back, corresponding to a second polymorph of symmetry Class 1. ASLTVS from Mcg crystallizes as antiparallel, anti-facial beta sheets packed face-to-face (Figure 6). This symmetry pattern is Class 5. Furthermore, its beta sheets are out-of-register (Figure 7A) because its strands are not perpendicular to the fibril axis, but instead are inclined. The third steric zipper, NFVFGT from Mcg displays a pattern of symmetry more complex than any previously observed steric zipper. The sheets that compose the ten previously described steric zipper symmetry classes can be built from translational repeats of one strand along the fibril axis (classes 1-4) or two strands (classes 5-10). Sheets of NFVFGT contain four orientations of the strand, and so require a minimum of four strands per translational repeat. The symmetry pattern of this sheet is one of six conceivable unique arrangements possible for a sheet with a four-strand translational repeat (Figure 7B).

**Figure 7.**
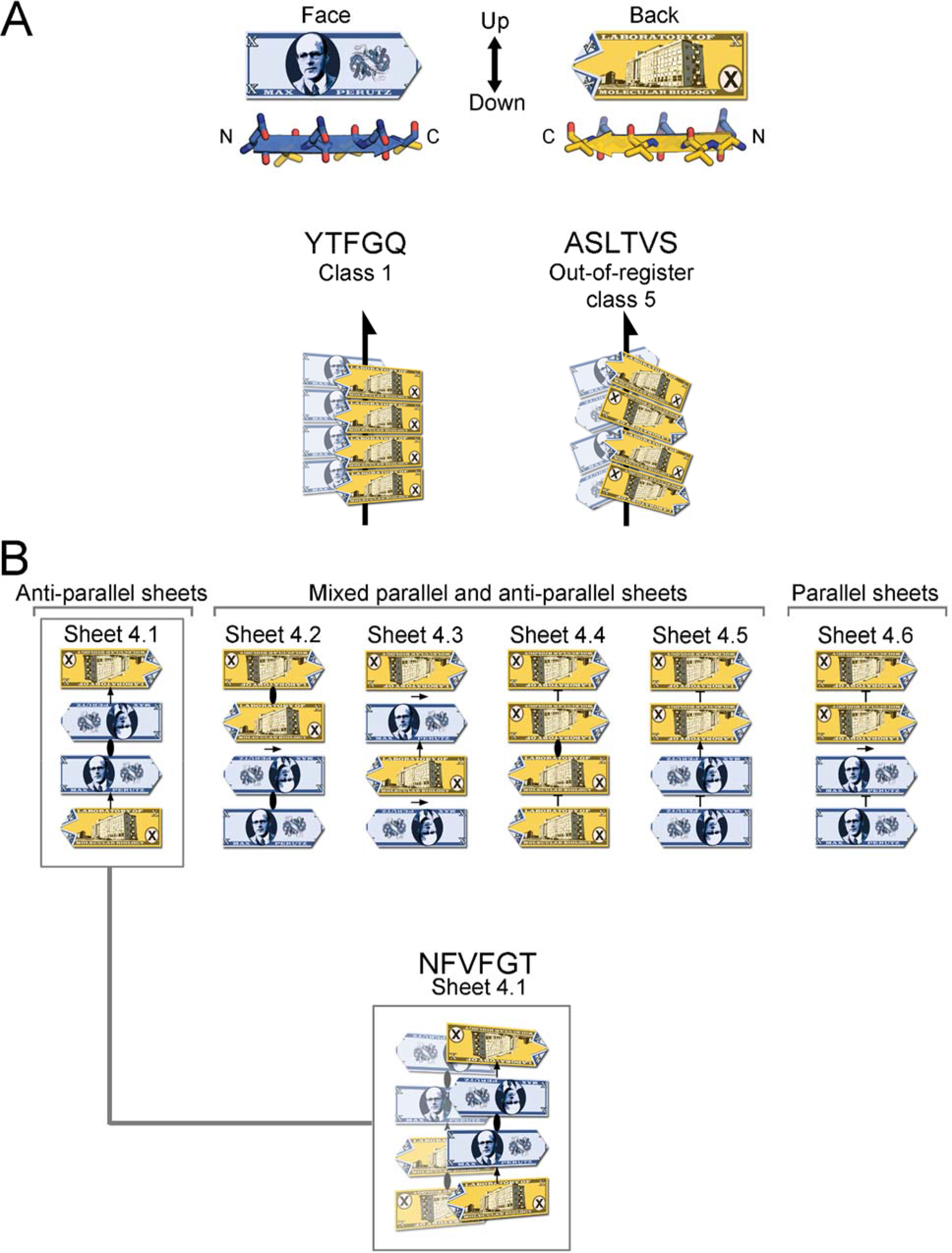
VL steric zipper and sheet geometry of amyloid spines, illustrated schematically with the fictitious Max Perutz banknote. **A.** The Max Perutz banknote represents a protein segment within a steric zipper; it has N- and C-termini, two distinct faces, and up- and down-direction of hydrogen bonds within a sheet. Arrows represent 2_1_ symmetry axes—meaning that the peptides are related by a 180° rotation about the arrow and a translated along the arrow of one-half the distance between Max Perutz banknotes. The YTFGQ steric zipper belongs to Class 1. The ASLTVS steric zipper belongs to Class 5. The packing of beta strands is out-of-register with its strands inclined relative to the fibril axis. **B.** The NFVFGT peptide crystallizes in a sheet containing four strands in the asymmetric unit. This sheet belongs to the one out of six conceivable arrangements of a sheet with a modulus of four strands and internal symmetry. Arrows represent two-fold symmetry operators, an ellipse represents a two-fold symmetry axis perpendicular to the page, and “T” represents translation.

## Discussion

Our results confirm that computational prediction of steric zipper propensity must be validated by experiments to correctly identify amyloidogenic segments. Computational prediction suggested several steric zippers within VLs, but site-directed mutagenesis verified only two such segments. We found that both are amyloidogenic independently of each other. Some of the other predicted high-propensity steric-zipper segments do indeed form amyloid fibrils, but only in isolation from the rest of the protein. That is, in the context of a full-length protein, not all predicted steric-zippers become exposed and available to drive formation of amyloid fibrils. Through a combination of computational, biochemical, and structural work, we have demonstrated the involvement of two distinct amyloidogenic segments in the formation of VL amyloid fibrils.

Our conclusions are consistent with previous solid-state Nuclear Magnetic Resonance (ssNMR) experiments. ssNMR experiments identified several segments having immobile conformations in VL amyloid fibrils, suggesting that some of these segments are involved in the formation of amyloid fibril spines. Piehl *et al.* and Hora *et al.* each assigned amino acids to these segments in the ssNMR spectra of amyloid fibrils of two VLs, human AL09 and murine MAK33(37–39). Two of the ssNMR-assigned segments correlate with our findings—they overlap with the amyloidogenic segments identified by our computational and site-directed mutagenesis experiments (Segment 1 in strand E of the Greek-key fold of immunoglobulin; and Segment 2 in strands F-G [Figure 8]). Although AL09 and MAK33 are different in their amino acid sequences and hence their amyloid fibrils are different, both include the amyloidogenic Segments 1 and 2. In our work, we have experimentally validated the involvement of Segments 1 and 2 in forming the steric zipper spine of the amyloid fibril. Thus, three different experiments, two ssNMR assignments of peptides in amyloid fibrils performed by independent laboratories and our site-directed mutagenesis, identify common regions involved in formation of VL amyloid fibrils (Figure 8A).

**Figure 8.**
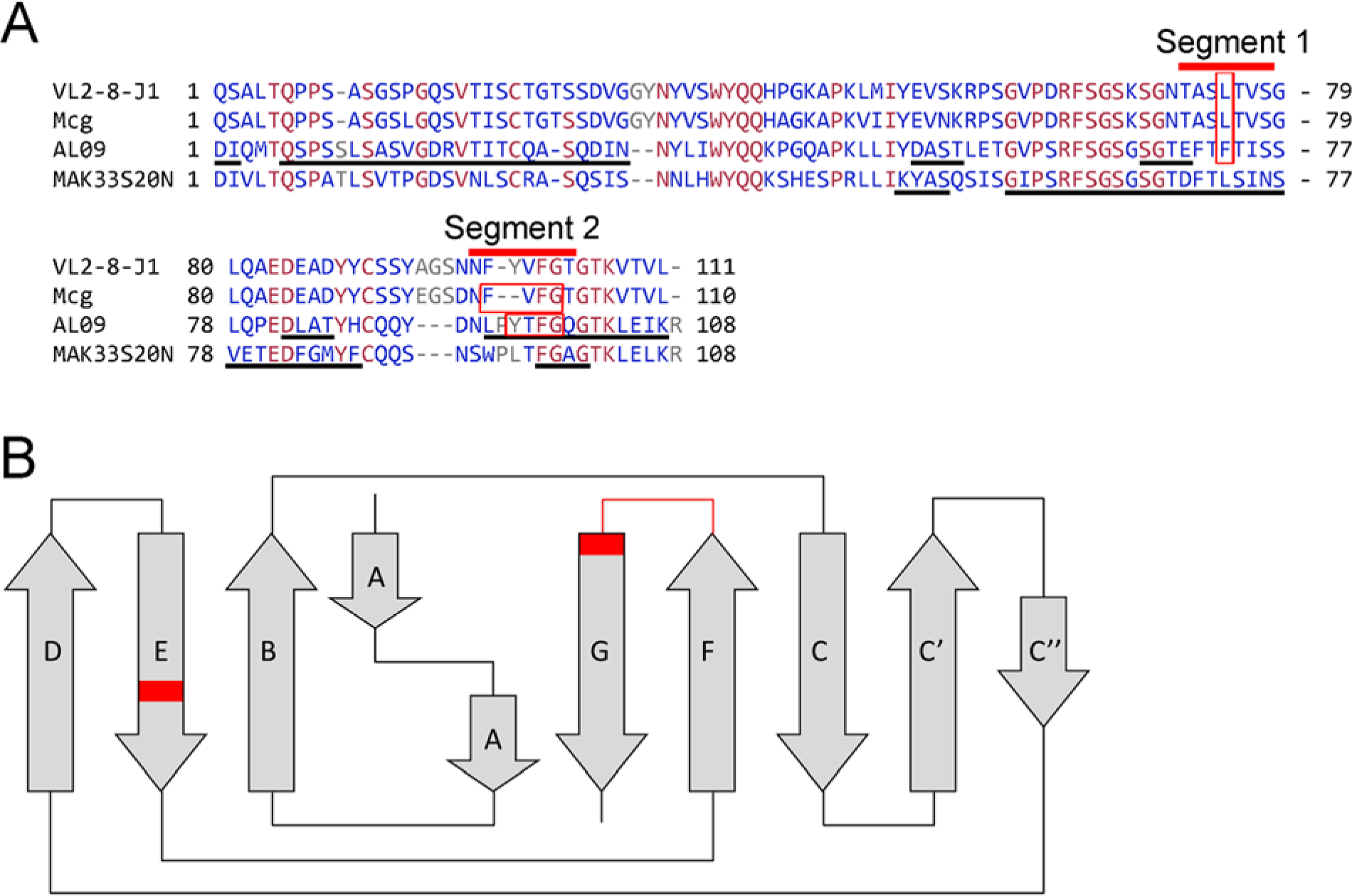
Sequence alignment of amyloidogenic VL Segments 1 and 2. **A.** Alignment of reference sequence VL2-8-J1, Mcg, Mak33S20N, and AL09. Underlined black lines denote segments assigned in amyloid fibrils by ssNMR experiments. Red bars identify amyloidogenic Segments 1 and 2. Residues in which mutations prevent formation of amyloid fibrils are enclosed in red boxes. Both Segments 1 and 2 overlap with MAK33S20N VL and AL09. Blue color indicates amino acids that vary among of the VLs. Red indicates conserved residues. **B.** The location of the amyloidogenic segments in the Greek-key fold of VLs, is indicated in red.

In support of our findings we succeeded in constructing geometrically-reasonable models of full-length VL amyloid fibrils with minimal deviation from crystallographic coordinates of the steric zippers (Figure 9). To minimize the degree of speculation in our models, residues outside the steric zipper segments maintain their native, globular fold, except for short regions that transition to the zipper spine. In doing so, the conserved disulfide bond within VL domains remains intact in all our models. Thus, the VL domains do not undergo complete unfolding. The models are free of serious steric clashes and preserve allowed Ramachandran geometry. As some measure of validation, we find that the diameters of the fibril models are consistent with the measured values for the diameters of amyloid fibrils in negative-stain electron micrographs, ^~^10-11 nm.

**Figure 9.**
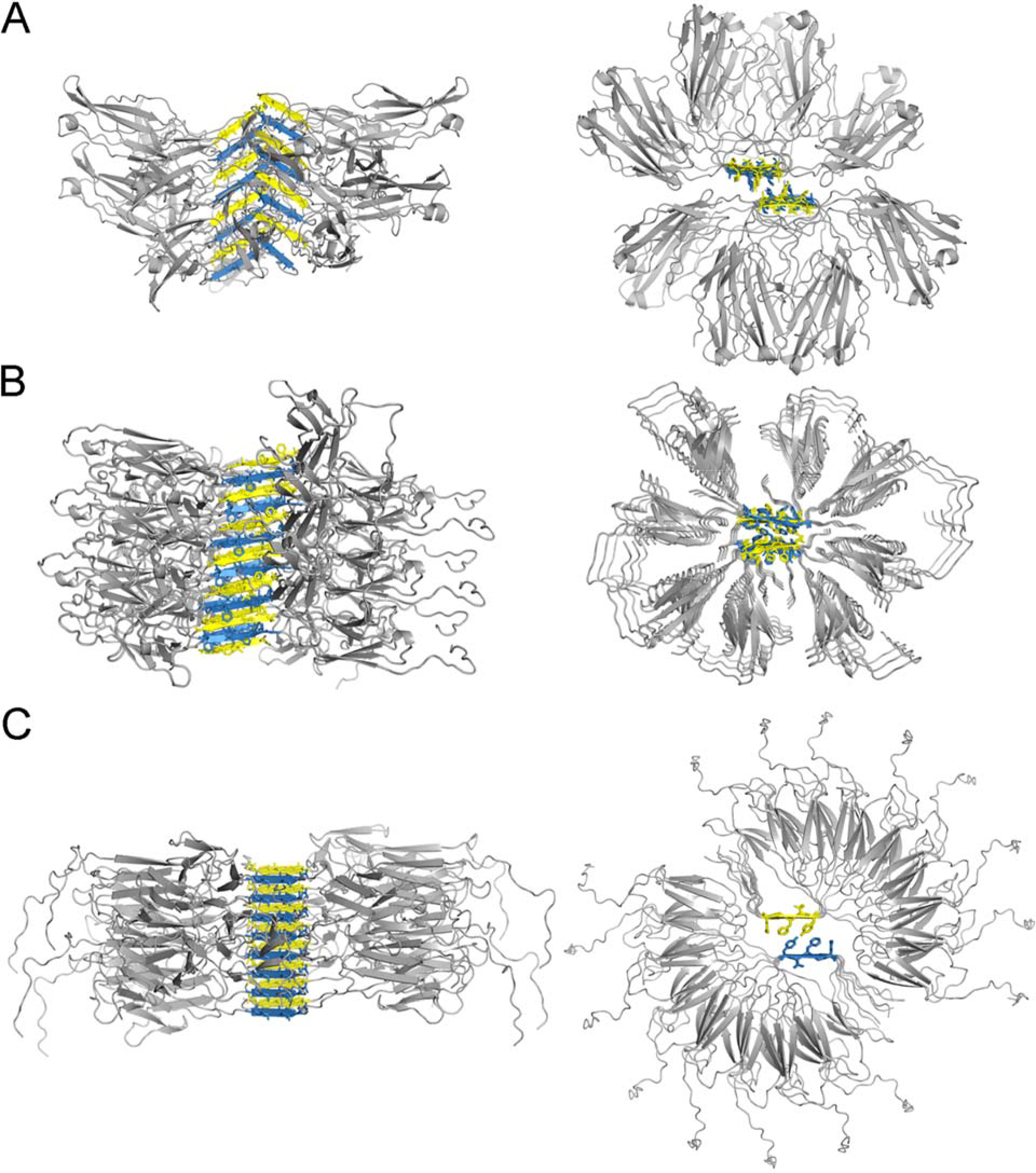
Speculative atomic models for VL amyloid fibrils based on the crystal structure of steric zipper spines. Each of the steric zipper spines forms when VLs partly unfold and expose the amyloidogenic segment to solvent. To model the full-length protein around the crystalline structures of the steric zipper spines, we used eight or 16 individual VL domains with somewhat different orientations. The left column shows models aligned sideways along the vertical axis of the fibril. The axes of fibrils propagate along the steric zipper spine through hydrogen bonding between the peptide backbones. The right panel shows the top view of the steric zipper spine and the interdigitating side chains. **A.** A model of the Mcg VL fibril based on the peptide derived from amyloidogenic Segment 1. **B.** A model of the Mcg VL fibril based on the peptide derived from amyloidogenic Segment 2. **C.** A model of the AL09 VL fibril based on the peptide derived from amyloidogenic Segment 2.

**Figure 10.**
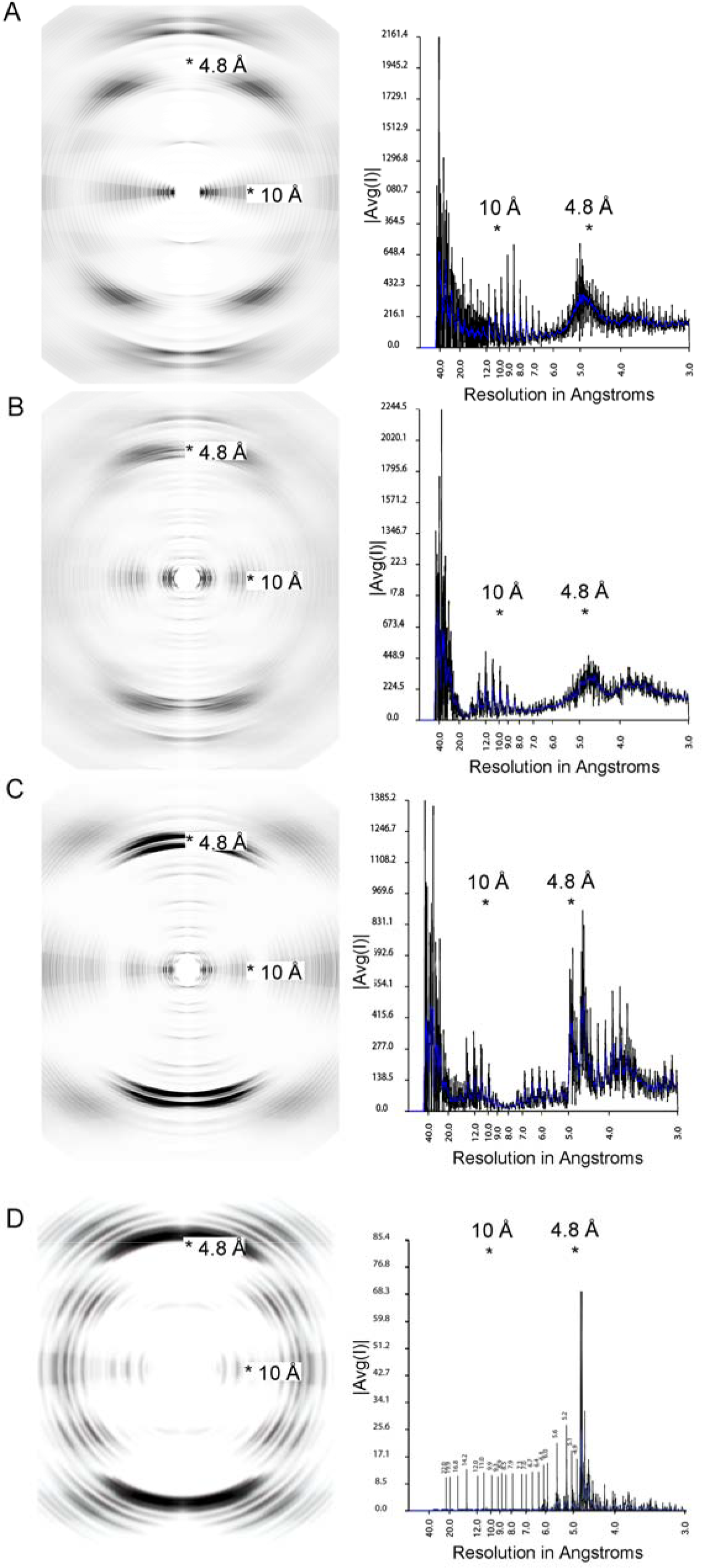
Fibril diffraction patterns (left column) and their radial integration profiles (right column) for amyloidogenic segments. The fibril diffraction patterns are calculated from the crystal structures of the segments (Figure 6). The radial integration profiles are calculated by cylindrical averaging of these single crystal diffraction patterns. These patterns show reinforcement of reflections in the vicinity of 4.8 and 10 Å, characteristic of steric zippers. In the right column, |Avg(I)| is the average intensity calculated from a Fourier transform of the atomic structures. **A.** Powder diffraction of ASLTVS. The meridional reflection near 4.8 Å is split due to inclined network of hydrogen bonds of the steric zipper spine. **B.** Powder diffraction of NFVFGT. **C.** Powder diffraciton of YTFGQ. **D.** Cylindrical averaging of single-crystal diffraction data of amyloidogenic segment EFTFTIS from kappa AL09. The averaged diffraction data of EFTFTIS shows strong reflections near 4.8 and 10 Å, perpendicular to each other. These reflections show the amyloid nature of the structure. The meridional reflection near 9.6 Å indicates that the structure contains an anti-parallel amyloid spine.

Identification of amyloidogenic segments has translational implications. Our conclusions directly bear on strategies for developing peptide-based inhibitors of VL amyloid fibrils. Current approaches to create peptide-based inhibitors generally aim to impede elongation of the amyloid fibril spine; modified peptides are designed to incorporate into the amyloid fibril spine with one face while arresting the propagation of the spine with the opposite face(40). However, in light of our results, it is clear that inhibitor design must simultaneously block at least two independent spines of VL amyloid fibrils. Future therapies may require a mixture of two or more inhibitors of amyloidogenic segments to halt fibril formation.

The structure of NFVFGT from the lambda VL Mcg reveals unprecedented complexity in its β-sheet geometry. It contains four distinct orientations of the β-strand within a single, out-of-register β-sheet. These features give the β-sheet a translational periodicity of 19.8 Å, which exceeds the more frequently observed in-register parallel sheets (4.8 Å), in-register antiparallel (9.6 Å), and out-of-register antiparallel (11.7 Å). Such lengthy periods, if truly present in biological fibrils, may produce more complex surface patches with affinity for more complex ligands than Thioflavin T or congo red, typically known to bind amyloid fibrils. It remains to be seen whether such complexity will be observed in other amyloid structures.

In summary, we experimentally identified two amyloidogenic segments within VLs. Each of the segments is a key propagator of amyloid fibrils, giving rise to distinct polymorphs. Both amyloidogenic segments occupy the same spatial location of the Greek-key immunoglobulin fold in both lambda and kappa VLs (Segment 1 in strand E of the Greek-key fold; and Segment 2 in strands F-G)(41). We found that this pattern of dual steric zippers is found in VLs belonging to both lambda and kappa classes, suggesting that homologous segments may be responsible for fibril formation in other VLs. Disrupting a single steric zipper is insufficient for blocking amyloid formation since an alternate zipper-forming region remains intact. These results explain previous difficulties in identifying a single residue or segment within VLs responsible for amyloid formation—there can be more than one. Our work offers an experimental framework to identify multiple amyloidogenic segments in VLs and other amyloid-forming proteins.

## Experimental procedures

### Computational analysis of VL sequences

Genomic and the variant sequences of VL2-8-J1 and Mcg were analyzed with ZipperDB (https://services.mbi.ucla.edu/zipperdb/) for steric zipper propensity, and with ConSurf (http://bental.tau.ac.il/new_ConSurfDB/) for amino acid conservation(32, 42).

### Preparation of recombinant Proteins

Protein samples were prepared as previously described. Site-directed mutagenesis was performed with PCR(43). DNA of every construct was verified twice by an external DNA-sequencing service (Laragen, Inc., www.laragen.com). In summary, VLs were expressed in *E. Coli*, purified in denaturing conditions, refolded, and concentrated(34). In the proline-scanning experiment, we found that four is the largest number of sequential prolines permits formation of the intended constructs. Longer proline segments resulted in degradation of several VL constructs and hampered the systematic experiments.

#### Amyloid fibril formation assays

Thioflavin (ThT) assays were performed as previously described(44). Assays were conducted in acidic conditions (50 mM NaAc/HAc pH 4 and 150 mM NaCl), 0.5 mg/ml of VL (40 µM), 37° C, and constant shaking at 300 rpm with Teflon balls with 0.125 inch radii as stirrers. ThT fluorescence spectra were recorded at 440/480 nm excitation/emission wavelengths. Each experiment was performed with freshly prepared samples in at least three independent biological repeats on different dates. Each experiment contained at least three technical repeats. Analysis of the genomic variant and its amyloid-stopping mutation (VL2-8-J1, VL2-8-J1-L75P) was performed in at least 14 different biological repeats.

### Electron microscopy (EM)

Samples from the ThT fibril formation assays were diluted with water to 10% v/v and applied onto copper grids with formvar-carbon coating (Ted Pella, Inc., Cat. No. 01810). Negative staining was performed with 2% w/v uranyl acetate and images were collected by means of a Tecnai T12 electron microscope at 120 kV with a Gatan CCD camera.

### Crystal structure determination

Peptides were purchased from Genscript, with trifluoroacetic acid salt substitution for HCl. The purity of peptides was greater than 95%. Peptides were dissolved in water to a concentration of 20 mg/ml and crystallization trials were set up using a Mosquito robot, with a 1:1 ratio of the peptide solution and crystallization screen. Crystals grew in hanging drops and were of needle-like morphology. Solutions that yielded the best-diffracting crystals are listed in Table 2. Crystals were dry-mounted onto pulled glass capillaries and diffraction data were collected at Advanced Photon Source beamlines 24-ID-C and 24-ID-E (ASLTVS—24-ID-E; NFVFGT—24-ID-C; YTFGQ—24-ID-C). Phases of were determined by molecular replacement with Phaser (ASLTVS) or by direct methods with Shelx(45, 46) (NFVFGT and YTFGQ). Molecular structures were refined with Refmac5 and deposited into the Protein Data Bank with codes 6DJ0 for ASLTVS, 6DIX for NFVFGT, 6DIY for YTFGQ(47).

**Table 2.** X-ray diffraction data and refinement statistics for the steric zippers that are responsible for amyloid fibril assembly by VLs.

|  | ASLTVS<br>PDB 6DJ0 | NFVFGT<br>PDB 6DIX | YTFGQ<br>PDB 6DIY |
| --- | --- | --- | --- |
| **Data collection** |  |  |  |
| Space group | P2<sub>1</sub> | P1 | C2 |
| Cell dimensions |  |  |  |
| *a*, *b*, *c* (Å) | 15.16, 11.58, 18.8 | 16.03, 11.66, 21.94 | 36.87, 4.81, 18.91 |
| $\alpha$, $\beta$, $\gamma$ (°) | 90, 97.32, 90 | 90.89, 103.23,<br>90.26 | 90, 104.95, 90 |
| Resolution (Å) | 18.6-1.3 | 20.0-1.0 | 18.3-0.9 |
| *R*<sub>sym</sub> or *R*<sub>merge</sub> (%) | 19.2 (27.0) * | 6.6 (12.4) | 6.2 (21.8) |
| *I*/σ*I* | 3.7 (1.7) | 12.8 (6.7) | 14.1 (3.2) |
| Completeness (%) | 80.5 (62.5) | 81.7 (62.0) | 90.7 (49.8) |
| Redundancy | 2.5 (1.5) | 3.4 (3.2) | 5.1 (3.0) |
| **Refinement** |  |  |  |
| Resolution (Å) | 18.64-1.3 | 21.3-1.0 | 18.3-0.9 |
| No. reflections | 1215 | 6154 | 2228 |
| *R*<sub>work</sub>/*R*<sub>free</sub> | 18.2/21.8 | 10.4/12.5 | 9.7/12.0 |
| No. atoms |  |  |  |
| Peptide | 93 | 388 | 82 |
| Water | 6 | 18 | 1 |
| B-factors |  |  |  |
| Peptide | 6.4 | 7.1 | 5.2 |
| Water | 12.8 | 21.2 | 23.2 |
| R.M.S deviations |  |  |  |
| Bond lengths (Å) | 0.019 | 0.015 | 0.013 |
| Bond angles (°) | 2.1 | 2.0 | 1.4 |
| **Crystallization conditions** |  |  |  |
|  | 0.1 M MMT (DL-malic acid:MES:Tris base; 1:2:2) pH 5, 25% w/v PEG 1500 | 0.1 M Imidazole HCl pH 8.0, 20% w/v PEG 3000, 0.2 M Zinc acetate | 2.8 M NaAcetate, 0.1 M Bis-Tris propane pH 7.0 |

Each structure was derived from a single crystal.
\*Highest resolution shell is shown in parentheses.

#### Construction of amyloid fibril models

In constructing our models of full-length VL amyloid fibrils we strived to make as few assumptions as possible. Therefore, atoms in the VL zipper spine were strongly restrained to maintain the arrangements observed in the crystal structures of their amyloidogenic segments. The remaining atoms of the VL were restrained to maintain the native globular folds (PDB codes 4UNU(34) and 3CDY(48)).

The largest challenge we faced in constructing the models was avoidance of steric clashes among globular domains, since the VL chains are very closely spaced along the fibril axis (4.8 Å for YTFGQ; 5.8 Å for ASLTVS; and 5.0 Å for NFVFGT). We performed a systematic search of orientation space, counting the number of steric clashes incurred for each possible orientation of the globular domains (in 5 degrees increments). To apply the incremental rotation operations and evaluate steric clashes, we used the programs PDBSET and CONTACT, respectively, from the CCP4 suite(49). We did not impose any twist in our fibril model, since there was no characteristic twist pitch length evident in electron micrographs of the fibrils. As a matter of convenience, we chose to model either eight or sixteen orientations of the globular domains around the fibril axis. We call this collection of unique orientations the asymmetric unit of the fibril. The fibril is then composed of translational repeats of this asymmetric unit (38.4 Å for YTFGQ, 19.8 Å for NFVFGT, and 46.3 Å for ASLTVS) along the fibril axis. For simplicity, the molecules within the asymmetric unit were restrained to have either cyclic or dihedral symmetry. Those models which produced the least number of clashes were manually edited with the program Coot(50), in order to build connections between the zipper spine and globular domain. The models were geometrically and energetically minimized with the programs CNS(51) and Rosetta(52).

## Acknowledgements

We thank Duilio Cascio, Daniel Anderson, and Michael Collazo for assistance with experiments. We acknowledge support from NIH NIA Grant 1R01AG048120-01 and the Howard Hughes Medical Institute. This work is based upon research conducted at the Northeastern Collaborative Access Team beamlines 24-ID-C and 24-ID-E, which are funded by the National Institute of General Medical Sciences from the National Institutes of Health (P41 GM103403). The Pilatus 6M detector on the 24-ID-C beam line is funded by a NIH-ORIP HEI grant (S10 RR029205). This research used resources at the Advanced Photon Source, a U.S. Department of Energy (DOE) Office of Science User Facility operated for the DOE Office of Science by Argonne National Laboratory under Contract No. DE-AC02-06CH11357.

## Author Contributions

B.B. devised the experimental plan and developed the project. B.B., S.R.E., and A.T.L, performed and analyzed the experiments. M.S. and B.B. built the atomic models of the amyloid fibrils. M.S., M.L., and B.B. performed the crystallography experiments. D.S.E. is the principal investigator overseeing the project. All authors participated in writing and commenting on the manuscript.

## Footnotes

* The content is solely the responsibility of the authors and does not necessarily represent the official views of the National Institutes of Health

